# Characterization Of Bitter Taste Receptor Dependent Autophagy in Oral Epithelial Cells

**DOI:** 10.1101/2024.02.02.578576

**Authors:** Nisha Singh, Saeid Ghavami, Prashen Chelikani

**Author notes:** Corresponding email address, D319, Department of Oral Biology, 780 Bannatyne Avenue, University of Manitoba, Winnipeg, MB R3E 0W4. Canada.

## Abstract

Microbial dysbiosis is an important trigger in the development of oral diseases. Oral keratinocytes or gingival epithelial cells (GECs) offer protection against various microbial insults. Recent studies suggest GECs expressed higher level of bitter taste receptor 14 (T2R14) compared to other taste receptors and toll-like receptors and acts as innate immune sentinels. Macroautophagy or autophagy is a cellular conserved process involved in the regulation of host innate immune responses against microbial infection. Here, we describe a robust method for evaluation of T2R14-dependent autophagy flux in GECs. Autophagy flux was detected using western blot analysis in GECs and further was confirmed using Acridine Orange dependent flow cytometry analysis.

**Workflow:** 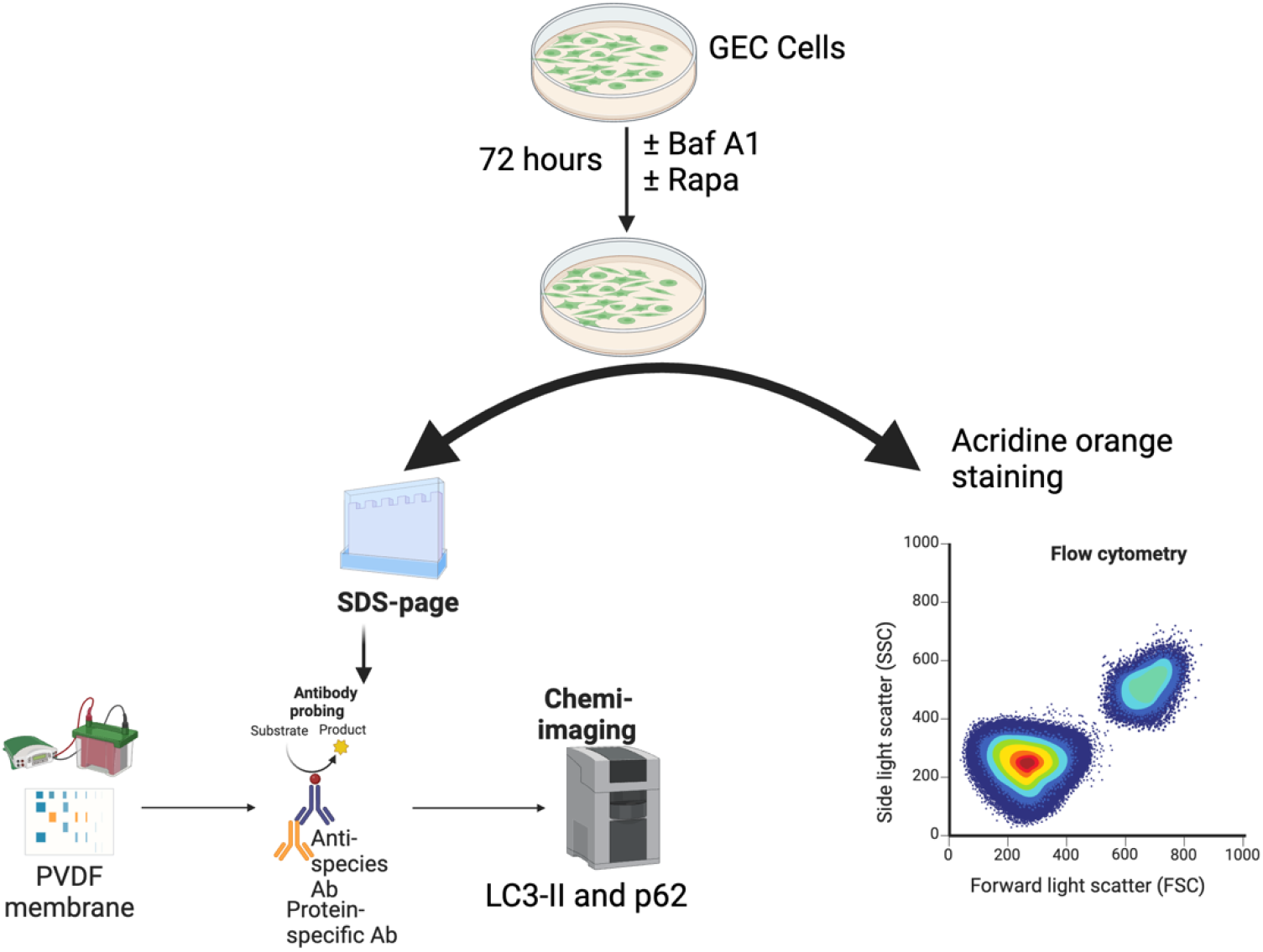

**Graphical abstract:** Schematic showing the methodology (Western blot and flow cytometry) used for assessment of autophagy flux in GEC (created with Biorender). GEC: Gingival epithelial cells, BafA1: Bafilomycin, Rapa: Rapamycin, LC3-II: microtubule associated light chain protein, p62: sequestosome 1

## 1 Introduction

Autophagy is a cellular process involved in breaking down macromolecules and organelles via the lysosomal degradation pathway (Germic, Frangez et al. 2019, Siapoush, Rezaei et al. 2023). Autophagy participates in the creation of autophagosomes, which transport cytoplasmic proteins and organelles to the lysosomal degradation machinery (Xie and Klionsky 2007, Alizadeh, da Silva Rosa et al. 2023). The formation of autophagosomes is regulated by key protein complexes such as mammalian target of rapamycin complex 1 (mTORC1) and Beclin 1. Conversely, AMP-dependent protein kinase (AMPK), a primary sensor of cellular energy levels, serves as a key activator of autophagy (Kim, Kundu et al. 2011).

In recent years, autophagy has emerged as a crucial regulator of innate immune functions which includes cytokine secretion, immune cell differentiation and clearance of pathogens (Deretic and Levine 2018, Deretic 2021). Previous investigations have well addressed the role of nutrient sensing G protein-coupled receptor (GPCR) in regulation of the autophagy pathway (Wauson, Zaganjor et al. 2012, O’Rourke, Kuballa et al. 2013). However, the involvement of majority of taste sensing GPCRs including the 25 T2Rs in the autophagy process is not well understood. Studies over the past decade suggest an important physiological and tissue specific role for T2Rs including host-microbe interactions and innate immune responses ((Medapati, Bhagirath et al. 2022). For example, activation of T2Rs resulted in increased nitric oxide (NO) production and enhanced phagocytosis of *Escherichia coli* and *Staphylococcus aureus* (Gopallawa, Freund et al. 2021). Our transcriptomic analysis of gingival epithelial cells (GECs) showed 21,448 genes and among them T2R14 or TAS2R14 was found to be expressed at high levels in different GECs (Medapati, Singh et al. 2021). The expression of TAS2R14 was notably elevated compared to the other 24 TAS2R genes (***p<0.001), as well as 8 TLR genes (***p<0.001).

Only few studies investigated the connection between T2Rs and autophagy (Pan, Sharma et al. 2017, Singh, Ulmer et al. 2024) Previous study in airway smooth muscles cells (ASMCs) suggested that bitter agonists quinine and chloroquine resulted in microtubule associated light chain protein (LC3β-II) accumulation and treatment of autophagy flux inhibitor bafilomycin A1 (Baf A1) and 3-methyl adenine (3-MA) leads to significant reduction of LC3β-II accumulation (Pan, Sharma et al. 2017). Oral keratinocyte or GECs produce many innate immune markers including nitric oxide and antimicrobial peptides (AMPs) against various oral pathogenic bacteria. Our recent investigation suggested that some cariogenic bacteria and their products show robust calcium responses in GECs and this effect is both T2R14 and autophagy protein 7 dependent (Singh, Ulmer et al. 2024). These results suggest that T2R14 is involved in determining the direction of autophagy flux in GECs via lysosomal degradation of autophagosome.

In a previous study, we reported that T2R14 modulates Gram-positive bacterial internalization (*Streptococcus mutans, Staphylococcus aureus*) in GECs (Medapati, Bhagirath et al. 2021). It is known that *S. aureus* utilizes the autophagosomes as a replicative niche (O’Keeffe, Wilk et al. 2015, Mulcahy, O’Brien et al. 2020). We showed that the internalization of *S. aureus* is T2R14 dependent in GECs, therefore it is important to analyze how cariogenic bacteria use the autophagy machinery for internalization and survival after evading the host immune system.

Here, we described two complementary methods for analyzing autophagy flux in GEC. The autophagic flux was characterized using western blot analysis of the autophagic markers SQSTM1 (encoding sequestosome 1 and is also known as p62) and LC3β-II (Dastghaib, Shojaei et al. 2020, Shojaei, Koleini et al. 2020, Alizadeh, Kochan et al. 2021). We also used Acridine Orange staining of acidic vesicular organelles (AVO) by flow cytometry to measure autophagy flux in GECs upon serum starvation, which is a physiological induction of autophagy (Hajiahmadi, Lorzadeh et al. 2023) (Singh, Ulmer et al. 2024).

## 2 Materials

### 2.1 Cell line

The oral gingival epithelial cell line GEC and the T2R14 knockout GEC cells were used in our previous studies (Medapati, Bhagirath et al. 2021, Medapati, Singh et al. 2021, Singh, Ulmer et al. 2024)

### 2.2. Cell culture and treatments

1. Keratinocyte growth medium-2 (KGM-2) for GEC cell culture was purchased from promo cell (Heidelberg, Germany Cat # C20011).
2. Acridine orange (CAS Number: 65-61-2), bafilomycin A1 (Cat#B1793-10UG) and rapamycin (53210-100UG) were purchased from Sigma-Aldrich Co. Canada.
3. Antibodies used in the study: The following antibodies were purchased: mouse SQSTM1/p62 antibody (5114) were purchased from cell singling (Canada Ontario). LC3β antibody (catalog number L7543) and monoclonal anti-β-actin (catalog number #A5441) were obtained from Sigma Aldrich in Oakville, Ontario, Canada. The goat anti-rabbit IgG-HRP conjugate (catalog number #17-6515) was sourced from Bio-Rad in Mississauga, Ontario, Canada, and the goat anti-mouse IgG-HRP conjugate (catalog number #A-10668) was also acquired from Bio-Rad in Mississauga, Ontario, Canada.
4. Flow cytometry

Beckman Coulter CytoFlex LX Digital Flow Cytometry Analyzer Four Laser System at the University of Manitoba, Flow cytometry Core facility.

## 3 Methods

### 3.1 Experiments for Western blot analysis

1. OKF6 cell both WT and T2R14 KO (40% confluent) were cultured in 10 cm tissue culture dishes (see **note 2**).
2. Treat the cells with media alone or media containing Bafilomycin (10 nM) and rapamycin (1000 nM) for 72 hours (see **note 2**).
3. After 72 hours, cell culture supernatants were collected in 15 ml tubes and cells were washed two times with 1X PBS. Finally, cell scraper was used to collect the GEC cells.
4. Cells containing media was centrifuged (350xg) and washed twice with phosphate buffer saline (PBS).
5. Cells were lysed in lysis buffer containing (1% Triton X 100 and Tris Buffer) using sonicator (3-5 pulses) and centrifugation at 10,000 g for 10 mins at 4°C to get rid of the cell debris. Supernatants was collected, and protein concentrations were determined using the DC protein assay (Bio-rad, Canada).
6. Cell lysates (10µg proteins loaded) for western blot for LC3β-II (1:2500) and p62 (1:1000), markers for autophagy flux were performed. 15% SDS -PAGE gels were run for separating the protein lysates.
7. Separated proteins were transferred to PVDF membrane for 90 mins at 4°C with the constant voltage of 100V, in blotting buffer (25 mM Tris, 192 mM Glycine and 20% Methanol).
8. Block the non-specific protein by blocking the PVDF membrane in 5% skim milk in Tris-HCl buffer (TBST) at room temperature with constant agitation for 1 hour.
9. Incubate the membrane with LC3β-II (1:2500) and p62 (1:1000) diluted in TBST buffer at 4°C with the constant agitation for overnight.
10. The following day, rinse the membrane three times for 10 minutes each with TBST buffer. Then, incubate the membrane with an HRP (Horseradish Peroxidase)-conjugated antibody (anti-mouse IgG, anti-rabbit IgG, or anti-goat IgG) diluted in TBST for 1 hour at room temperature.
11. Reveal the antibody reaction by chemiluminescence using ECL Plus and normal ECL (Bio-Rad, Canada) (see **note 4**).
12. Incubate the blots sequentially at room temp for 2 hr with monoclonal anti-β-actin antibody (Sigma) diluted at 1:10,000 and process as described above.
13. Standardize the densitometry readings of positive bands by comparing them to the corresponding value of β-actin. Chemiluminescence was assessed using the Bio-Rad ChemiDoc MP imaging system, and the blot densities were analyzed using Image Lab version 5.2.1 software from Bio-Rad.
14. Statistical significance was calculated using Two-way ANOVA with Tukey’s post-hoc test (see **note 1**).

### 3.2 Experiments with Acridine Orange (AO) Flow cytometry analysis

1. OKF6 cell both WT and T2R14 KO (20,000) were seeded in 12 well plates under serum starved conditions (see note 1).
2. Treat the cells with media alone or media containing Baf A1 (10nM) and rapamycin (1000 nM) for 72 hours (see **note 3**).
3. After, 72 hours, cells were trypsinized with 0.25% Trypsin-EDTA (Invitrogen, Canada) for 10 mins at 37°C. Cells were trypsin neutralized with DMEM F12 media containing 10% serum and centrifuged for 4 mins at 300 g to get the cell pellets (see **note 5**).
4. OKF6 cells were stained with acridine orange (final concentration of 1µg/ml) and incubated at 37°C for 10 mins in dark (see **note 6**).
5. Stained cells were washed once with PBS, and the flow cytometry analysis using cytoflex flow cytometer was performed for evaluating the autophagy flux (Red and Green fluorescence) (Thome, Filippi-Chiela et al. 2016, Hajiahmadi, Lorzadeh et al. 2023) (see **note 7**).
6. Acquisition was done by using channels in the red (PECy5) and green (FITC) lasers and recording 10000 events.
7. Acridine orange flow plots were analyzed on Flow jo Version 10 software (TreeStar, Ashland, OR)

## 4 Results

### 4.1 Western blot analysis of autophagy flux in GEC

Our previous studies suggest T2R14 is primarily involved in mediation of innate immune responses to bacterial infection in GECs (Medapati, Bhagirath et al. 2021, Medapati, Singh et al. 2021). To determine whether and how oral bacteria utilize the autophagic machinery of GEC, it is necessary to characterize the role of T2R14 in modulating autophagic flux (Singh, Ulmer et al. 2024).

To assess the impact of T2R14 on autophagy flux in GECs, oral keratinocyte cells (OKF6 Wt for wild type and OKF6 T2R14 KO for knockout) were employed as previously described (Medapati, Bhagirath et al. 2021). Autophagic flux was assessed via western blot analysis of the autophagic markers p62 and LC3β-II. Both cell lines underwent treatment with an autophagy flux inhibitor, Bafilomycin A1 (Baf A1, 10 nM), and an autophagy flux inducer, rapamycin (1000 nM), for 72 hours. This treatment of GECs did not result in any noteworthy alteration in cell viability. After 72 hours of treatment, cells were lysed and the LC3 lipidation and p62 degradation was analyzed by western blot. These results suggested that in the basal condition (i.e. without Baf A1 or Rapa treatment), more LC3β-II (^*^p<0.05) and p62 accumulated in T2R14 KO cells relative to WT, suggesting that fewer autophagosomes were being degraded in the T2R14 KO GECs (Fig. 1 A-C). To further confirm this, both cells were treated with autophagy flux inhibitor Baf A1 for 72 hours. Baf A1 treatment leads to further increase in LC3β-II and p62 accumulation in both WT and T2R14KO GECs (Fig. 1 A-C). Treatment of GECs with Baf A1 blocked lysosomal degradation of autophagosome efficiently and thereby led to an accumulation of autophagic vesicles. Next, the cells were treated with rapamycin, an autophagic flux inducer, and changes in LC3β-II and p62 were assessed. LC3β-II and p62 accumulation was reduced in T2R14 KO cells relative to WT under conditions of induced autophagic flux (Fig. 1 A-C). Overall, on the basis of LC3β-II and p62, the presence of T2R14 in GECs downregulates autophagy flux under basal conditions.

**Figure 1.**
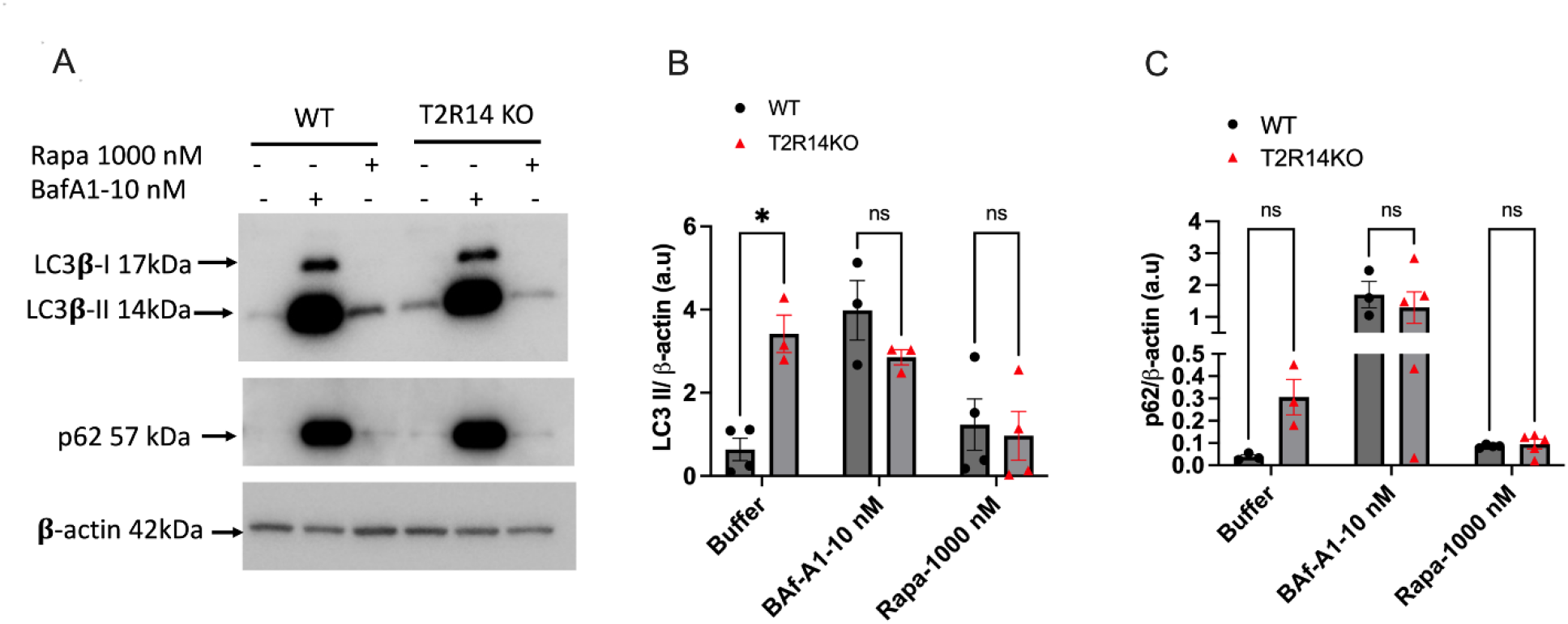
T2R14 mediated autophagy flux in GEC. A. Western blot analysis of autophagy flux in GEC. **A**. Western blot detection of LC3β-II and p62 autophagy markers in WT and T2R14KO cells upon incubation with autophagy blocker bafilomycin (Baf A1-10 nM) and activator rapamycin (Rapa 1000 nM) for 72 hours. **B and C**. Quantification of relative levels of LC3β-II/actin and p62/actin by densitometric analysis. The bar graphs were generated using graph pad prism 9.0. Statistical significance was calculated using Two-way ANOVA with Tukey’s post-hoc test ^*^p<0.05, ^**^p<0.01. The data for B and C represents SEM of 3-4 independent experiments.

### 4.2 Acridine Orange staining for detecting acidic vesicular organelles (AVO) by flow cytometry analysis in GEC

Acridine Orange staining is a quick and robust assay for monitoring and detecting AVO (Thome, Filippi-Chiela et al. 2016, Klionsky, Abdel-Aziz et al. 2021). WT and T2R14KO GECs cell lines were serum starved for 72 hours to induce the autophagy (Ghavami, Yeganeh et al. 2018, Shojaei, Koleini et al. 2020). Cells were additionally subjected to treatment with Baf A1 (10 nM) and Rapa (1000 nM) for 72 hours to assess autophagy flux (measured through red and green fluorescence) using a CytoFLEX flow cytometer (Thome, Filippi-Chiela et al. 2016). Our findings indicate that under conditions of nutrient deprivation, which promote autophagy, T2R14KO cells exhibit a significant rise in red fluorescence compared to WT cells, while no change in green fluorescence is noted (Fig. 2). Moreover, treatment of WT cells with the autophagy flux inhibitor Baf A1 results in a complete loss of red fluorescence (Fig. 2). Notably, BafA1 treatment inhibits vacuolar H(+)-ATPase (V-ATPase), preventing lysosomal acidification and consequently eliminating red fluorescence. Intriguingly, the autophagy flux inducer Rapa and serum starvation lead to an increase in red fluorescence, indicating that stimuli inducing autophagy boost the number of acidic vesicular organelles (AVO) (Fig. 2). Hence, based on AVO, the presence of T2R14 in GECs influences the abundance of actively acidic lysosomes.

**Figure 2.**
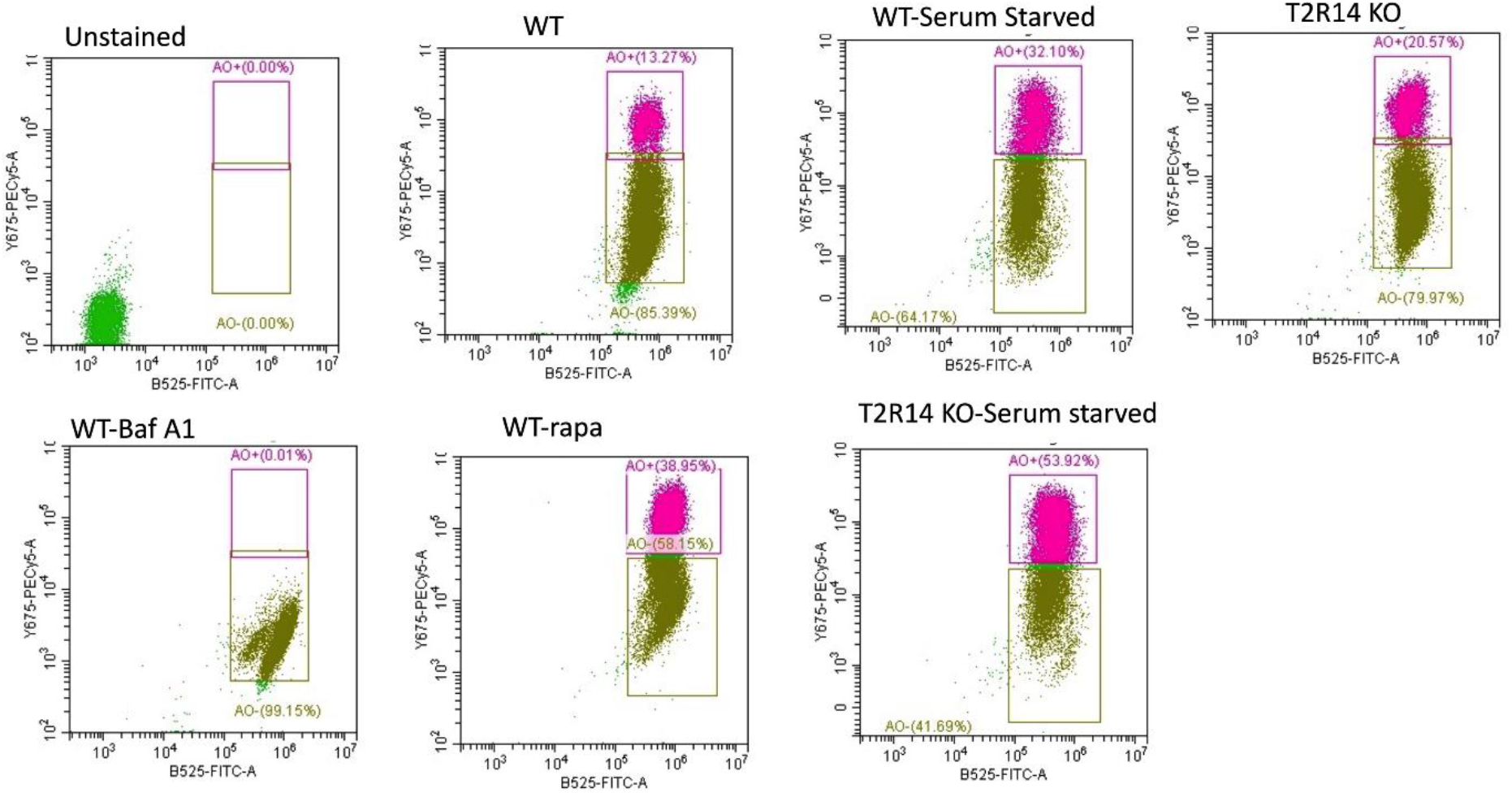
Acridine Orange staining for detecting acidic vesicular organelles (AVO) by flow cytometry analysis in GEC. Flow cytometry plots depicting red, green, and the red/green fluorescence ratio were utilized to assess autophagy flux in both WT and T2R14KO GECs under serum-starved conditions and normal growth conditions. Representative flow plots illustrate the acquisition of data on red and green fluorescence in acridine orange-stained GECs (WT and T2R14 KO) subjected to serum starvation or treated with Baf A1 (10 nM) and Rapa (1000 nM) for 72 hours to detect autophagy flux using red and green lasers. Acridine orange flow plots were analyzed using FlowJo Version 10 software (TreeStar, Ashland, OR).

## 5 Notes

1. It is recommended to have at least three replicates for each condition, although having a minimum of three replicates is necessary for statistical analysis. Ideally, it would be beneficial to conduct four or more replicates for each condition. Additionally, ensure that cells are serum-starved and grown in 1% growth factor as specified in the Acridine Orange staining protocol
2. OKF6 cell culture media preparation and cultures should be performed in a Biosafety cabinet.
3. After exposure to Baf A1 and rapa treatments monitor the OKF6 cell morphology under the light microscope for 72 hours.
4. Avoid long exposure of blots for LC3β-II for over exposure of bands especially in Baf A1 and rapa treatments.
5. OKF6 cells are very adherent cells, so use 0.25% trypsin EDTA for cells trypsinization and immediately neutralize the cells with DMEM F12 with 10% serum containing media and spin down the cells at 300xg for 5 mins.
6. The tubes containing the acridine orange dye were placed in a rack and covered with foil paper because acridine orange is light sensitive and can easily degrade.
7. Acridine Orange based autophagy flux flow cytometry should be performed within 30 mins of staining.

## Acknowledgement

This work was supported by Grant No. PJT-159731 from the Canadian Institute of Health Research (CIHR) to PC.

## Author Contributions

N.S conducted the experiments and analyzed the data. N.S and P.C wrote the manuscript. P.C; N.S and S.G contributed to the conceptualization, design and data interpretation. All authors critically revised the manuscript.

